# Laboratory efficacy of a solar-powered spatial repellent device against *Anopheles* mosquitoes

**DOI:** 10.64898/2026.08.13.744622

**Authors:** Robert T. Jones, Jessica Dennehy, Matthew A. Turner, Will Dyall, Freya I. Spencer, Isaac Owusu, Akin Jenkins, Alexandra Hiscox, Sarah Y. Dewhirst, James G. Logan

## Abstract

Spatial repellents represent a promising approach to complement existing malaria vector control interventions by reducing contact between mosquitoes and humans. However, the efficacy of passive spatial repellent devices may be affected by environmental conditions, particularly temperature and airflow, which can influence the release of volatile active ingredients. Active-release devices may provide more consistent delivery of spatial repellent compounds. We investigated the efficacy of a commercially available battery-powered spatial repellent device and a low-power solar-powered device designed for potential use in Africa. Laboratory trials were conducted to evaluate the efficacy of the two spatial repellent devices against *Anopheles* mosquitoes. Protective efficacy was assessed by comparing mosquito probing on human participants during spatial-repellent and control tests. The effect of the devices on mosquito entry into the test chamber was also assessed. The protective efficacy was 86% with the commercial battery-powered device and 80% with the low-power solar-powered device. A Wilcoxon rank-sum test showed that there was no significant difference between the performance of the two devices in terms of protective efficacy (p = 0.856) or entry inhibition (p = 0.7989). The low-power device could be charged using a small, household-level photovoltaic panel, to provide a potentially practical means of delivering spatial repellents in off-grid settings.

## Introduction

Strategies to control malaria and other vector-borne diseases in Africa rely heavily on the use of insecticide treated nets and indoor spraying of residual insecticides. However, repellent technologies represent another fundamental aspect of preventing mosquito-borne disease transmission, and remain relatively underused.^1^

Topical repellents are available with both natural and synthetic active ingredients. Whilst some have been shown to provide many hours of protection,^2^ they require high user compliance and there remains insufficient evidence that topical repellents can prevent malaria in settings where other vector control interventions are in place.^3,4^ By contrast, spatial repellents represent an exciting alternative approach to malaria prevention by deterring host-seeking female mosquitoes from taking blood meals without the need for frequent reapplication of formulations to human skin.^5^ Further, they are not burdened with the high implementation costs and logistical deployment obstacles that are associated with alternative technologies, such as the development of sterile strains of mosquitoes and mass-trapping techniques. Instead, programmes can readily deploy spatial repellents using current technologies and at relatively little cost.

Several active ingredients have been shown to provide spatial repellent effects and provide protection from mosquito bites.^6^ These have been incorporated into devices with a range of formats. Although few studies have investigated effects on epidemiological outcomes, two studies of limited power suggest that the volatile pyrethroids transfluthrin and metofluthrin, when used as coils inside people’s houses, have potential uses as part of malaria control programmes.^7,8^ A disadvantage of such devices is the necessary production of smoke inside the home, which may cause adverse effects after extended exposure.^9^

These compounds have been incorporated into passive devices that do not utilise external electrical heating or combustion. Transfluthrin-treated eave ribbons offered significant protection against mosquitoes in hut trials,^10^ as well as in small field trials.^11^ Such devices can be situated to allow for the gradual release of active ingredient through natural airflow, but given that they rely on passive emanation, low temperatures are expected to hinder the vaporization of the transfluthrin and reduce their efficacy in cooler evenings and in colder climates. Investigations with transfluthrin-treated hessian strips found that preceding-day temperatures affected the number of bites received by mosquitoes at night. Although there was >90% protection against *Anopheles* bites on nights when the preceding daily mean temperatures were 23°C or greater, protection was approximately 80% on nights with preceding daily mean temperatures below 21°C.^12^

The first results of more recent trials with plastic film transfluthrin-based devices that are applied onto interior walls have demonstrated that passive devices can reduce the risk of malaria infection in an African setting characterised by high malaria transmission.^13^ Participants in a trial in Kenya reported that, unlike bed nets that only provide protection at night, the device protects against mosquito bites in the evening, although some also raised concerns about the consistency of the product’s effectiveness over time, suggesting that the amount of active ingredient declined with continued use.^14^ Together, these observations suggest that maintaining consistent airborne concentrations of repellent compounds may be challenging under field conditions. Active release could help to stabilise delivery and ensure consistent outputs at biologically significant concentrations despite fluctuations in airflow and temperature.

Active release may be achieved through internal exothermic reactions, which raise the local temperature within a device to promote evaporation.^15^ Electrically-powered spatial repellent devices are also available in some countries for those travelling to regions where they might be bitten by mosquitoes. These devices do not produce smoke but instead repel mosquitoes by the slow release and circulation of active ingredients from a quiet, battery-powered fan. Here, we investigated the efficacy of a commercially available battery-powered repellent device in a laboratory trial. The study informed the development of a low-power alternative device that may be made available to users in off-grid settings, with the electricity required for the fan being produced from a small, household-level solar panel. This low-power device was also tested in a laboratory trial.

## Methods

### Experiment 1: Mosquito repellent laboratory study with commercial device

#### Test devices

The commercial battery-powered device (Mijia Smart Mosquito Repellent) was manufactured by Xiaomi (Beijing, China). It was supplied with 2 x AA batteries to accompany each transfluthrin-treated cartridge.

#### Insects

*Anopheles coluzzii* (Coetzee & Wilkerson, 2013) were obtained from reference strains held at the London School of Hygiene & Tropical Medicine (LSHTM). The mosquitoes were reared and housed under optimal environmental conditions of 25°C±2°C and 80% relative humidity (RH) with a 12:12 h light:dark photoperiod. Non-blood fed female mosquitoes aged 3-5 days, starved of sugar, were used in the trial.

#### Ethical approval

The study was given a favourable opinion by the LSHTM Research Ethics Committee (application number 22624). All participants recruited to the study provided informed written consent. All participants were provided training in mosquito collection prior to taking part and had a bite test to assess their reaction to *An. coluzzii*.

#### Testing procedure

Repellent devices were tested in tiled chambers at LSHTM. The arena consisted of two adjoining chambers (one ‘test’ and one ‘release’ chamber), each with a capacity of approximately 10 m^3^, connected by a single door. The temperature and humidity were maintained at 27 ± 2°C and 75 ± 10% RH.

A control repellent device with no active ingredient was placed into a corner of the ‘test’ chamber, approximately 1.5 m from a centrally positioned stool, and the connecting door was closed. A participant entered the test chamber 1 h later, wearing a bee suit, shoe covers and gloves for protection. The participant sat on the stool and rolled their sleeves to the elbow to expose their lower arms. The connecting door was left open. A batch of 50 mosquitoes was then released from the adjoining ‘release’ chamber. Over a 15-minute period, any mosquitoes that probed on the participant’s exposed arms were collected by mechanical aspirator. After the 15-minute test period, the door connecting the two chambers was closed, and any uncollected mosquitoes were captured. The number of mosquitoes collected (1) probing, (2) in the test chamber, and (3) in the release chamber, was counted. The two chambers were washed with 70% isopropyl alcohol and left to aerate before further tests were conducted.

The test was repeated with a repellent device containing the active ingredient. The device was activated 1 h prior to the test to allow for release of transfluthrin into the test chamber. Following the test, the repellent device was left switched on in a large plastic container for 168 h to simulate use 12 h per day for 14 days. The devices were checked at 1-3 day intervals to ensure they were still running and to ventilate them. After the aging period, the device was tested again following the same procedure. Testing was repeated with a new devices until four participants had completed testing.

#### Mortality assays

A control device, without repellent cartridge, was placed in one corner of the test chamber and the connecting door to the release chamber was closed. The device was left for 1 h. After this time, the door was briefly opened and a batch of 50 female mosquitoes was released into the test chamber. The connecting door to the release chamber was then closed. After 15 minutes, the door was opened and the mosquitoes collected by mouth aspirator.

Following the control test, an unused repellent device containing the active ingredient was switched on and placed in the corner of the test chamber. The connecting door to the release chamber was closed. It was left for 1 h. After this period, a batch of 50 female mosquitoes was released and re-captured as above.

All recovered mosquitoes were transferred into a paper cup covered with a mesh and supplied with cotton wool soaked in 10% glucose solution. The cup was labelled and placed in a Perspex recovery chamber at 27 ± 2°C and 75% ± 10% RH, and observed for knockdown after 1 h and mortality after 24 h. Mosquitoes that were moribund or dead were classified and recorded as knocked down at 1 h and as dead at 24 h.^16^

#### Air sampling

To quantify the concentration of transfluthrin in the air, air sampling was conducted using an AirLite pump (SKC, Blandford Forum, Dorset, UK) and Tenax TA thermal desorption tubes (Part No: C1-AAXX-5003) (Markes International, Bridgend, South Wales, UK) in a subset of the trials. Samples were taken 1 h after the control device had been switched on (n = 4), immediately before the participant entered the chamber, and 1 h after the transfluthrin device had been switched on (n = 5). The sampler was placed at ground level approximately 1 m from the spatial emanator device. Air was sampled over 2 minutes. The pumps were calibrated with a TD tube and flow meter to 0.5 L/min. The mass of transfluthrin collected was determined using gas chromatography–mass spectrometry (GC-MS).

The GC-MS analysis was performed using a 7890B gas chromatograph and 5977B MSD mass spectrometer (Agilent Technologies, Cheshire, UK) equipped with a Unity Xr thermal desorption sampler (Markes International , Bridgend, South Wales, UK). Samples were desorbed using a two stage desorption method where Initial sample desorption temperature was 300 °C and was held for a time of 5 min. The desorbed sample was transferred a Tenax TA cold trap held at 5 °C. The final trap desorption was performed at 300 °C and held for 5 minutes. The Thermal desorption tranferline temperature was set to 200 °C. Chromatographic separation of transfluthrin was performed using at Rxi-5 MS column, (30 m, 0.25 mm, 0.25 µm) (Restek Thames, Buckinghamshire, UK). Column temperature was held at 60 °C for 3 min and increased to 180 °C at 10°C/ min and finally increased to 300 °C at 20 °C. The total run time was 30 minutes. The carrier gas was helium and the flow rate was set to 1.5 ml /min.

The temperature of the Ion source and transfer line were 150 °C and 310 °C respectively. Detection of transfluthrin was carried out in a the SIM /SCAN mode where a mass range of 40-400 amu for scan component of the method. SIM conditions contained three ions m/z 91, 163 and 335, which are indicative of tranfluthrin. All data was collected at a scan rate of 6.6 scans/s. Transfluthrin had an approximate retention time of 19.4 minutes and was quantified using m/z 163 with comparison against a single point started prepared from a certified tranfluthrin reference material (Dr Ehrenstorfer, LGC, Surrey, UK) at a concentration of 1 ng on column mass.

#### Statistical analysis

Protective efficacy (PE) was determined as a proportion by comparing the number of mosquitoes probing on the participants in the spatial-repellent tests to the control tests. PE = ((C_l_-T_l_)/C_l_)*100 where C_l_ is the number of mosquitoes probing on the participant during control testing and T_l_ is the number of mosquitoes probing on the participant during testing of the spatial repellent device.

The reduction in vector entry into the test chamber between spatial-repellent tests and controls tests was determined as a proportion. Vector entry inhibition = ((C_e_-T_e_)/C_e_)*100, where C_e_ is the number of mosquitoes that entered the test chamber during testing of the control device and T_e_is the number of mosquitoes that entered the test chamber during testing with the spatial repellent device.

### Experiment 2: Mosquito repellent laboratory study with solar-powered device

#### Test devices

The solar-powered device (Mossie-GO) was developed and manufactured by Africa Power Ltd (Cowfold, England). These were provided with a transfluthrin-treated cartridge. The device requires 0.205W to operate the fan, and can be charged using a 5W panel. They were charged for the purposes of this study by a micro-USB cable.

#### Insects

*Anopheles gambiae* (Giles, 1900), strain G3, were obtained from reference strains held at Digital Odour Technologies Ltd. The mosquitoes were reared and housed under optimal environmental conditions of 27°C ±2°C and 75% ±10% RH with a 12: 12 h photoperiod. Non-blood fed female mosquitoes aged 5-7 days, starved of sugar, were used in the trial.

#### Testing procedure

The tests were performed using the same procedures as described in Experiment 1, in chambers of approximately 15 m^3^ at Digital Odour Technologies Ltd. Participants in the study were members of the study team blinded to the treatment/control device being used. All participants were provided training in mosquito collection prior to taking part and had a bite test to assess their reaction to the mosquito species. Testing was repeated with new devices and refill units until four participants had completed testing.

#### Air sampling

Air sampling was performed as described above, but with the pumps running for 15 minutes. Samples were collected from both control and treatment tests.

#### Statistical analysis

Analyses of protective efficacy and entry inhibition were performed as for Experiment 1. The PE and entry inhibition of the two devices were compared using Wilcoxon rank-sum tests. Analyses were performed in RStudio (R-4.6.0).^17^

## Results

### Experiment 1: Mosquito repellent laboratory study with commercial device

#### Protective efficacy

PE of the commercial battery-powered device against *An. coluzzii* was 86% prior to aging and 94% after 2 weeks of use. At the same time points, entry inhibition was determined to be 10% and -23%, respectively.

#### Mortality assays

None of the mosquitoes (0/50) recovered from the test chamber after exposure to the control device were determined to be knocked down at 1 h (0% knocked down), and none were dead at 24 h (0% mortality). Of the 50 mosquitoes released into the test chamber with the device containing the active ingredient, 46 were dead or moribund after 1 h (92% knocked down). After 24 h, 41 were dead (82% mortality).

#### Air sampling

The mean mass of transfluthrin collected 1 h after switching on the battery-powered device with a transfluthrin cartridge was 2.14 ng (1.39 SD), which corresponds to an air concentration of 2.1 ng/L. The mean mass of transfluthrin collected 1 h after switching on the control device was 0.08 ng (0.09 SD), or approximately 0.1 ng/L (Figure 1).

**Figure 1.**
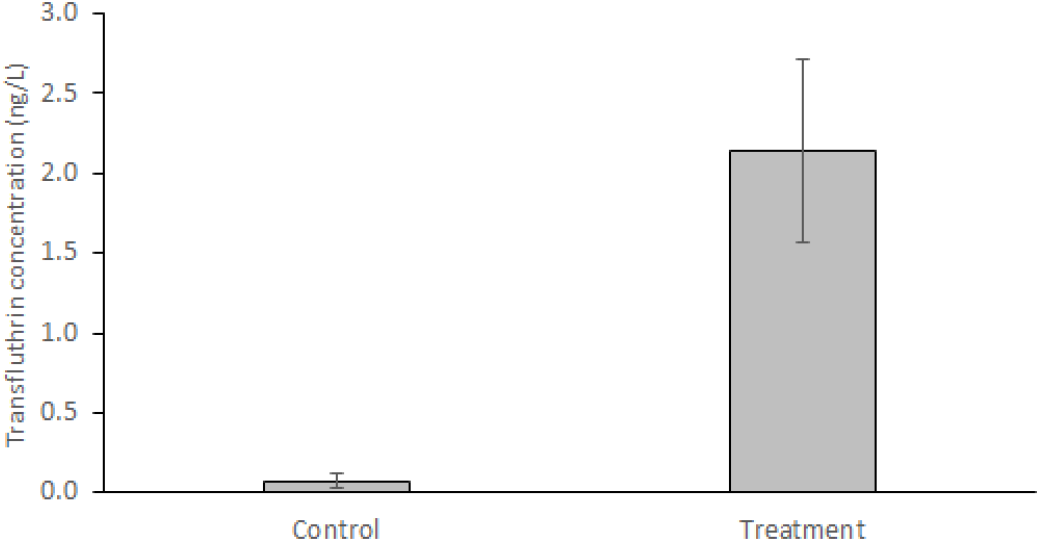
Mean (with standard error bars) concentration of transfluthrin detected in control (n=4) and treatment (n=5) tests, 1 h after the device was switched on. The test chamber had a volume of approximately 10 m^3^.

**Figure 2.**
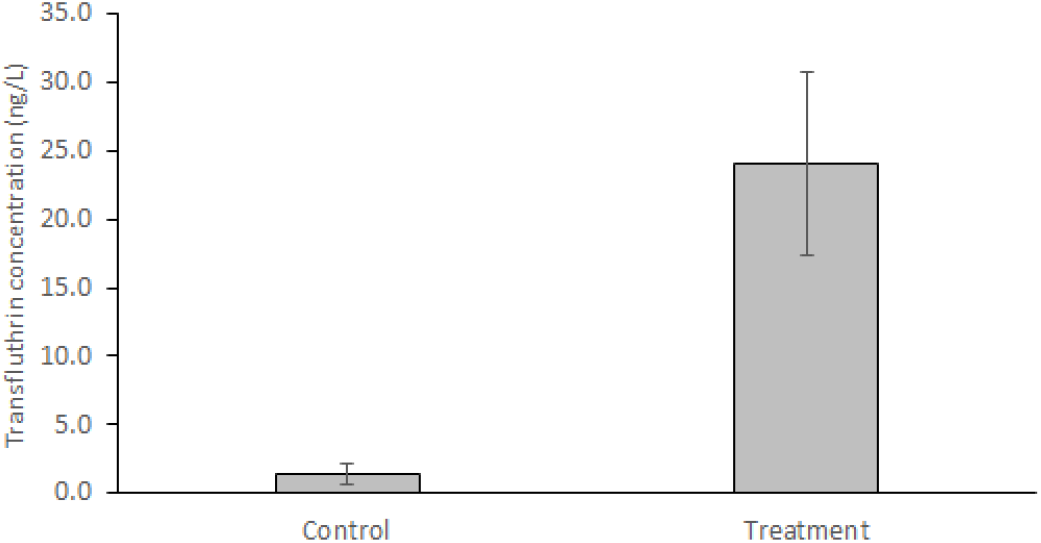
Mean (with standard error bars) concentration of transfluthrin in control (n = 2) and treatment (n = 4) tests, 1 h after the device was switched on. The test chamber had a volume of approximately 15 m^3^.

### Experiment 2: Mosquito repellent laboratory study with solar-powered device

#### Protective efficacy

PE of the solar-powered device against *An. gambiae* was 80%. Entry inhibition was determined to be 10%. Wilcoxon rank-sum tests showed that there was no significant difference between the performance of the two devices (PE: p = 0.856; entry inhibition: p = 0.7989).

#### Air sampling

The mean mass of transfluthrin collected 1 h after switching on the battery-powered device with a transfluthrin cartridge was 180.15 ng (SD 100.28), which corresponds to an air concentration of 24.02 ng/L. The mean mass of transfluthrin collected 1 h after switching on the control device was 10.5 ng (SD 7.78), or approximately 1.4 ng/L (Figure 1).

## Discussion

The plateau in the progress towards malaria elimination has stimulated interest in novel interventions that might supplement insecticide treated nets and indoor residual spraying.^18^ The trials described here with a commercial battery-powered device provided a benchmark for the development of a low-powered alternative that could be charged by a small photovoltaic panel. Such a device could offer protection through powered spatial emanation that has not previously been available in off-grid settings.

We found that the Mossie-GO device matched the performance of the commercial device in terms of protection from bites, despite requiring much less power to operate. This indicates that it achieved comparable entomological protection with a lower energy requirement. In practical terms, this could permit the use of electrically-powered spatial repellent devices in off-grid settings where their use has previously not been viable.

While the present study did not find evidence of reductions in mosquito entry into protected spaces, deterrence is just one feature of the mosquito behaviours collectively referred to as spatial repellency.^19^ Anti-feeding, reduced fecundity, knockdown and mortality have been reported in response to exposure to transfluthrin and affect the entomological parameters of malaria transmission.^20–23^ Vector bites are critical for disease transmission, and along with daily mosquito mortality, are the most important parameters for the determination of disease risk.^24^ The most direct indicator of the potential value of a spatial repellent is, therefore, its impact on host-seeking behaviour, measured here and elsewhere using human landing catches.

Human landing catches provide an approximation of the number of potentially-infectious mosquitoes that could bite one person at a particular time and place, and offer a suitable measure of protective efficacy while limiting the risk of vector-borne disease transmission to study participants by interrupting mosquitoes before they have taken a blood meal.^25,26^ While differing experimental setups have been used, human landing catch-based estimates of protective efficacy have been reported for transfluthrin-based devices in a range of formats,^27–33^ and are typically determined as a primary entomological endpoint prior to the evaluation of novel spatial repellent interventions under field conditions. Despite their utility, recent modelling has indicated that the additional effects of some tools, such as the lethal effects of volatile pyrethroids, mean that reductions in landing can be underestimated by human landing catch methods. As a result, human landing catches tend to under-predict the relative reduction in vectorial capacity in susceptible mosquito populations. ^34^

The airborne concentrations of transfluthrin required to elicit biological responses in susceptible mosquitoes have been investigated in previous studies. Repellency and other sub-lethal effects can occur at concentrations lower than those required for toxicity, but exact airborne concentrations have been difficult to determine because they often fall below the detection limits of available analytical methods.^35^ Rigorous analyses correlating mosquito entry into spaces protected by spatial repellents with the chemical concentration in the air remain limited.^36^ Measuring responses of *Anopheles* to transfluthrin along a gradient from the exposure source, Martin et al. reported that where transfluthrin concentration could be detected (corresponding to at least ∼0.32 ng/L), there was a 1.6-fold higher knockdown/mortality compared to locations where transfluthrin was below this detection limit.^37^ However, repellency was similar across the gradient, and no clear association between repellency and measured transfluthrin concentrations above versus below the detection threshold could be identified. A more complete understanding of the airborne concentrations at which spatial repellents elicit behavioural and toxic effects would help to inform modelling studies and target product profiles. In particular, defining the concentrations that drive repellency over space and time would support optimisation of delivery formats and deployment strategies,^36^ including the frequency at which devices need to be replaced, how many units are required in houses of different sizes, and the extent to which ventilation and absorptive materials within homes limit the availability of active ingredients in the air.

We recognise the limitations of the present study. The devices were tested against colony-reared mosquitoes and were used in the controlled settings of indoor arenas. Different species of *Anopheles* were used in the two experiments because of availability at the time of the trials. We were only able to conduct limited air sampling in the indoor arenas and, given the increased dispersion of the Mossie-GO compared to the other commercial device, the air sampling results can only be treated at an estimate as they fall out of the linear dynamic range of the analytical method. Further, we have not investigated how natural airflow might influence indoor transfluthrin concentrations. However, assessments of protective efficacy provided by the Mossie-GO device in semi-field experimental huts have recently been completed, and the results from those studies and those presented here suggest that the device could be used, alongside existing interventions, to contribute to reductions in malaria transmission by inhibiting human-vector contact. This will be explored through a field trial with epidemiological outcomes.^38^

## Acknowledgements

We thank the study participants, and the research ethics committee for their review of our protocols.

